# Mating-Type Imputation (MTI) provides an efficient tool for the mating-type inference of tetrapolar fungi

**DOI:** 10.1101/2025.09.01.673225

**Authors:** Haixu Liu, Yu Wang, Ruiheng Yang, Weijun Li, Huiyang Xiong, Yu Li, Yongping Fu, Shijun Xiao, Dapeng Bao

## Abstract

Mating-type identification is fundamental for the genetic diversity and genetic breeding study in fungi, especially for tetrapolar basidiomycetes, whose mating-types are determined by two multiallelic loci A and B. Traditional mating-type identification of monokaryons relies on manual inference based on their hybridization experiments; however, the process is rather complex, time-consuming and error-prone for large-scale hybridization experiments. In this study, we developed the Mating-Type Imputation (MTI) software for automatic, rapid and accurate monokaryon mating-type inference for tetrapolar fungi using a combinatorial pruning traversal algorithm. Taking the compatibility matrix from 435 hybridization experiments of 30 *Flammulina velutipes* monokaryons as an example, MTI takes a few minutes to infer the accurate mating-types of all monokaryons, which might take several days for manual inference by experienced investigators. Furthermore, MTI enables us to investigate how false-positives and false-negatives influence mating-type inference results. Using simulated compatibility matrix, we found that MTI could accurately detect potential false-negatives in compatibility and successfully infer the true mating-type combination with limited false-negatives, but the tool could be misled by any false-positives to result in a wrong mating-type combination, indicating that false-positives records in hybridization experiments should be strictly obviated for mating-type inferences. In summary, MTI provides an efficient tool for mating-type inference of tetrapolar fungi, offering technical support for the research paradigm in mating-type studies of edible and medicinal fungi and holding significant theoretical value and broad application prospects in the fields of fungal genetic diversity and breeding studies.

## 1. Introduction

Mating-type research is the hot-spot direction in mycological genetics and evolutionary biology, facilitating our understanding of fungal species isolation, genetic diversity, population structure, adaptive evolution, and cross breeding(Fraser & Heitman, 2003; Kinoshita et al., 2018). Mating-types determine the mode of sexual reproduction in fungi. Research indicates that mating-types control the fuse of gametes and the formation of sexual spores, affecting the genetic diversity and the population phenotypic variation(Nieuwenhuis et al., 2011). Meanwhile, the highly diversified multiallelic mating-type loci in fungi provided valuable information for fungal taxonomy and germplasm tracing studies(Kinoshita et al., 2018; Ryan & Smith, 2004). Previous studies have shown that the varieties of mating-types could reflect the population diversity of fungi and the mating-type alleles can serve as reliable molecular markers for taxonomic units clarification and strain identification. From an applied perspective, understanding mating-types holds significant guiding value for fungal breeding and germplasm utilization. In the breeding of economically important fungi such as medicinal and edible fungi, knowledge of parental mating-types is fundamental for controlling hybridization direction and improving the efficiency of genetic improvement. Therefore, fungal mating-type research not only elucidates the fundamental principles of fungi at the genetic and evolutionary levels but also provides theoretical support for applied research in taxonomy and breeding investigations(Bao & Wang, 2016; Kothe, 2001; L. Larraya et al., 1999).

The tetrapolar mating system is a prevalent, complex and highly evolved sex recognition system in basidiomycetes(Freihorst et al., 2016; Kües, 2015). This system operates via two mating-type loci, commonly referred to as the A locus and the B locus, respectively controlling hyphal gamete recognition (A locus, encoding transcription factors such as HD1/HD2) and fusion behavior (B locus, encoding peptide pheromones and receptors)(Raudaskoski & Kothe, 2010). Most commercially cultivated edible and medicinal fungi, such as *Flammulina velutipes*, *Lentinula edodes*, *Ganoderma sichuanense*, and *Auricularia auricula*, exhibit a typical tetrapolar mating system(H. Li et al., 2022; Wang et al., 2016; Yu et al., 2022). Identification of the mating-type A/B locus of parents is essential for designing monokaryon hybridization, enabling purposeful recombination of desirable traits and enhancing breeding efficiency(L. M. Larraya et al., 2001; Y. Zhang et al., 2019). Additionally, in agricultural and ecological systems, the prevalence of diseases caused by tetrapolar basidiomycetes (such as rusts and smuts) is closely related to their mating-type distribution(Kües, 2000; F. Liu et al., 2021; Xu et al., 2017). Therefore, accurate mating-type identification is crucial for understanding disease transmission mechanisms and formulating control strategies(Argiroff et al., 2023; Rokas, 2022).

Strain compatibility matrix deduced from pairwise monokaryon hybridization experiments is the main information for mating-type identification of tetrapolar basidiomycetes, but the manual mating-type inference process is complex, time-consuming and error-prone(Dong et al., 2022). Especially when the population size grows, the workload for manual mating-types inference increases exponentially. Although many studies of mating-type identification have been reported before, mainly on edible and medicinal mushrooms, a universal, efficient and automated mating-type inference method based on strain compatibility matrix is still lacking for tetrapolar fungi (Kothe, 2001; Krogerus et al., 2021; W. Li et al., 2024). Consequently, it is an urgent need in current fungal genetics research to develop such a mating-type inference tool using compatibility information from hybridization experiments for large-scale monokaryons, which will not only improve the efficiency and accuracy of mating-type identification but also provide strong technical support for research in fungal population genetics, evolutionary biology and breeding(Sun et al., 2019).

Based on these considerations, this study developed a software, Mating-Type Imputation (MTI), to infer monokaryon mating-types using monokaryon hybridization experiments for tetrapolar basidiomycetes. Taking advantage of the high rate of compatibility among monokaryons, a combinatorial pruning traversal algorithm is applied to improve the efficiency of mating-type inference for large-scale monokaryon hybridization experiments. MTI was validated in the mating-type research of *Flammulina velutipes* and used to investigate how errors in the strain compatibility matrix influence the results of mating-type inference. In summary, MTI and its application in this study not only provides a reliable tool for the mating-types inference of tetrapolar basidiomycetes, but also helps to establish the paradigm for the monokaryon mating-type studies in general fungi.

## 2. Materials and Methods

### 2.1 Development and application of mating-type inference method

#### 2.1.1 Mating-type inference (MTI) algorithm

To achieve accurate mating-type inference, MTI software acquires three inputs (**Figure 1**). First, the parental information of the monokaryon strains. Second, the strain compatibility matrix deduced by clamp connection observations in hybridization experiments(Bao et al., 2004; Raudaskoski, 2015; S.-S. Zhang et al., 2023). Third, the loci consistency table deduced by colony morphology observations for incompatible hybridization combinations in OWE-SOJ experiments(Fangcan et al., 2000).

**Figure 1:**
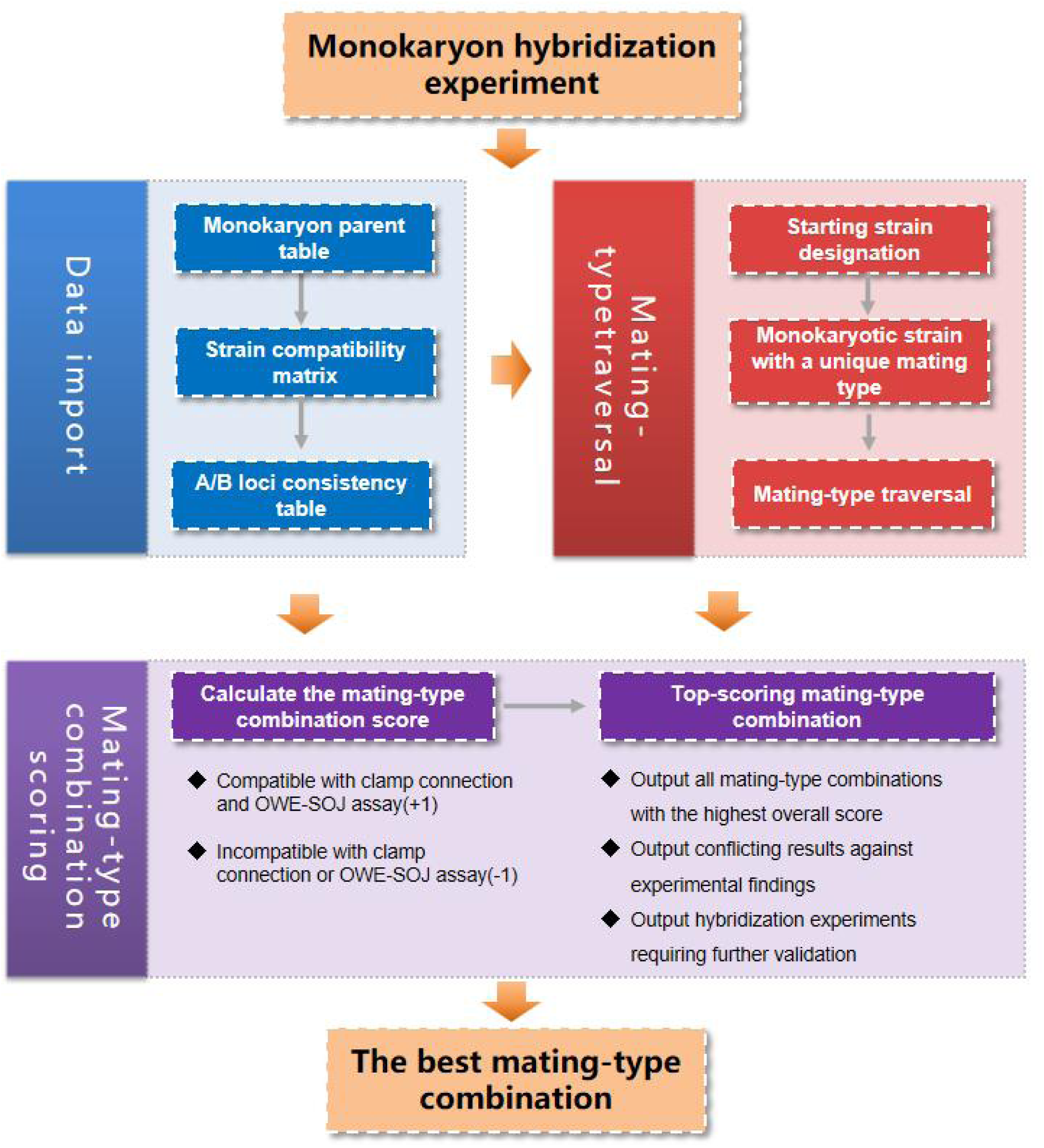
Analysis process of MTI mating-type inference software.

The workflow of MTI for mating-type inference follows the below steps (**Figure 1** and **Figure 2**). Initially, users are required to provide two monokaryon strains with clearly distinct mating-types as the starting strains, typically using two monokaryon strains derived from one dikaryotic parent, with their mating-types designated as A_1_B_1_ and A_2_B_2_, respectively. Next, a sub-population of monokaryon strains exhibiting pairwise compatibility in the hybridizations are selected and those monokaryons were labeled as A_3_B_3_, A_4_B_4_ and so on. Subsequently, the mating-types of the remaining monokaryons, showing any incompatibility in pairwise hybridizations, were traversed to generate all possible mating-type combinations. Then, scores of all mating-type combinations were calculated to represent their overall consistency with the compatibility matrix (**Figure 1**). Finally, the mating-type combination with the highest score is identified as the best mating-type combination based on the hybridization experiments.

**Figure 2:**
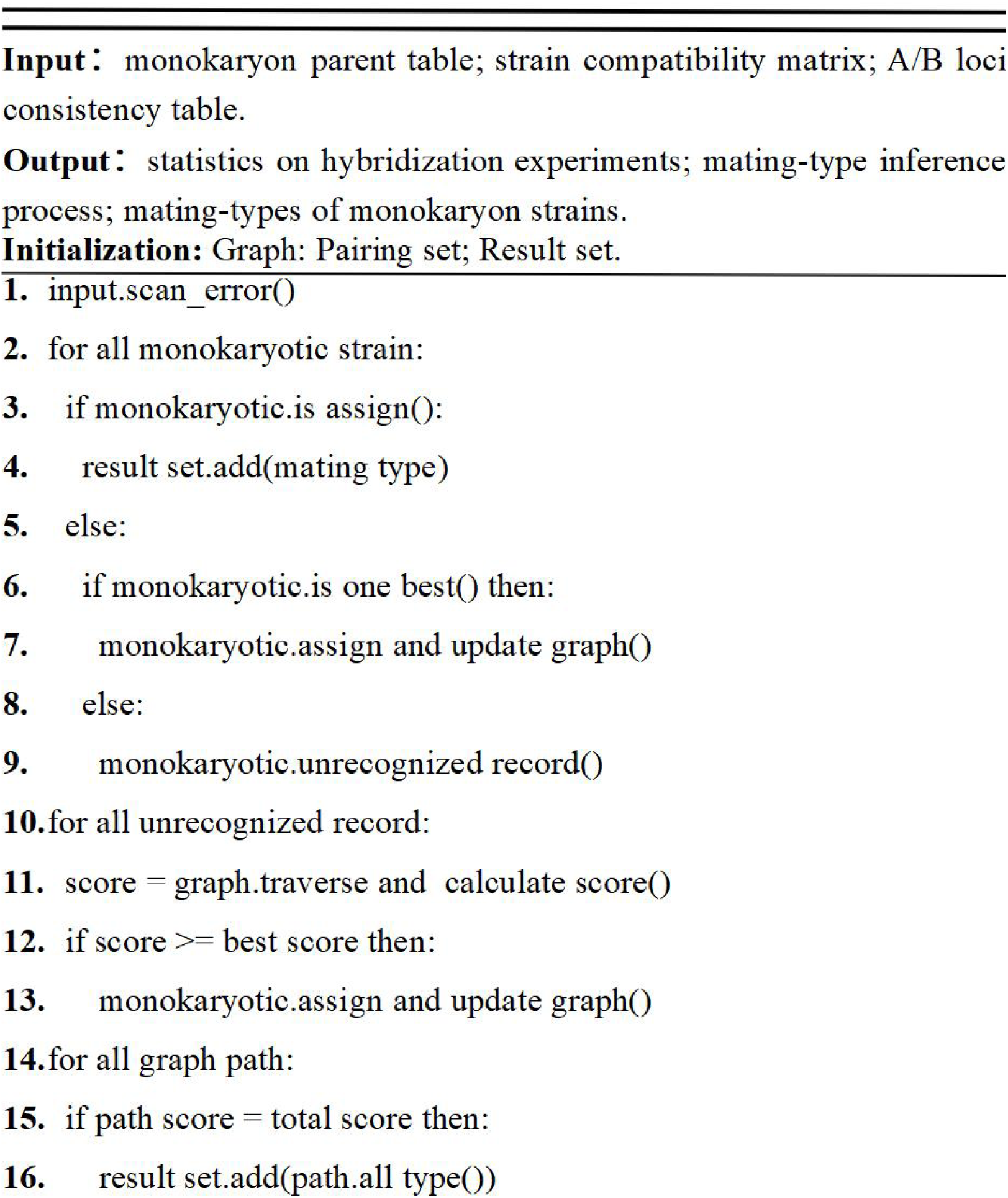
Pseudo code of MTI for mating-type inference algorithm.

#### 2.1.2 Online deployment of MTI

The MTI platform is developed using TypeScript and Kotlin. The web front-end is developed with a declarative and component-based technology model based on the Vue front-end framework, which effectively reduces page refresh time, lowers maintenance difficulty, and enhances user experience. The server back-end is developed using the Ktor asynchronous framework, which offers advantages such as rapid application startup, low performance overhead, and high security, significantly improving data analysis speed(Adamczyk et al., 2011). For the visualization of mating-types and sample data, the open-source OpenTiny HUICharts package is employed. This package is a version based on ECharts and features advanced charting capabilities and rapid responsiveness(W. Li et al., 2022; Nebeling & Norrie, 2013). For the optimal experience, it is recommended to access the website using JavaScript-supported browsers such as Mozilla Firefox and Google Chrome.

### 2.2 Monokaryon hybridization experiment of *Flammulina velutipes*

#### 2.2.1 Strain information

The 14 wild strains and 1 cultivated *Flammulina velutipes strain* used in this experiment were provided by the Edible Fungi Research Institute of the Shanghai Academy of Agricultural Sciences (**Table 1**). Through the technique of protoplast monokaryons separation(Bao et al., 2020; Chang et al., 1985; Rodriguez-Iglesias & Schmoll, 2015), 30 monokaryon strains were obtained.

**Table 1:** Information on the binucleate strains used in this study.

| ID | Strain number | Collection site | Source |
| --- | --- | --- | --- |
| 1 | GS0002 | Guan'e Gou Deadwood, Longnan, Gansu Province | Wild type |
| 2 | GS0116 | Ping Shan Tang Dong Road, Yangzhou City,<br>Jiangsu Province | Wild type |
| 3 | GS0145 | Bengxing Village, Guihu Town, Chaozhou City,<br>Guangdong Province | Wild type |
| 4 | GS0146 | Dong Cao Wa, Guqiao Bridge, Fengtai County,<br>Huainan City, Anhui Province | Wild type |
| 5 | GS0169 | Campus of Huazhong Agricultural University,<br>Wuhan City, Hubei Province | Wild type |
| 6 | GS0174 | Huayuan Street, Xinjin District, Chengdu City,<br>Sichuan Province | Wild type |
| 7 | GS0175 | Yangxunqiao Street, Keqiao District, Shaoxing City,<br>Zhejiang Province | Wild type |
| 8 | GS0180 | Longtian Township, Yongxin County, Ji'an City,<br>Jiangxi Province | Wild type |
| 9 | GS0193 | Zhuzang Town, Zhijin County, Bijie City, Guizhou<br>Province | Wild type |
| 10 | GS0194 | Liufangtou, Gaoshan Town, Dongkou County,<br>Shaoyang City, Hunan Province | Wild type |
| 11 | GS0196 | Shaugulin, Qianjiang Town, Qianjiang District,<br>Chongqing Municipality | Wild type |
| 12 | GS0200 | Fufei Wubu, Fuzhou City, Fujian Province | Wild type |
| 13 | GS0216 | Huancheng Road, Xiangcheng City, Zhoukou City,<br>Henan Province | Wild type |
| 14 | GS0225 | Lv Yinhu Street, Duyun City, Qiannan Buyei and<br>Miao Autonomous Prefecture, Guizhou Province | Wild type |
| 15 | T011 | Nagano Prefecture, Japan | Cultivation type |

#### 2.2.2 Culture media preparation

Potato Dextrose Agar (PDA) medium was prepared by dissolving 39 g of PDA powder in distilled water and bringing the final volume to 1 L(Y. Liu et al., 2002). The solution was sterilized by autoclaving at 121°C for 20 minutes.

Oak Wood Extract (OWE) medium was prepared using the following method(Fangcan et al., 2000): 50 g of oak wood chips were suspended in distilled water, and the volume was adjusted to 1000 mL. The suspension was boiled for 60 minutes and subsequently filtered through double layers of gauze. Separately, 20 g of agar was dissolved in distilled water and the volume was brought to 1,000 mL. A 200 mL aliquot of the oak wood filtrate and the entire 1,000 mL agar solution were sterilized independently by autoclaving at 121°C for 20 minutes. After cooling both components to approximately 50°C, they were thoroughly mixed and poured into Petri dishes to solidify as plates.

Squeezed Orange Juice (SOJ) medium was prepared by first dissolving 20 g of agar in distilled water and bringing the final volume to 1,000 mL (Fangcan et al., 2000). This agar solution was combined with 200 mL of freshly squeezed orange juice. Both components were sterilized independently by autoclaving at 121°C for 20 minutes.

After cooling to approximately 50°C, the sterilized agar solution and orange juice were thoroughly mixed and poured into Petri dishes to solidify as plates.

#### 2.2.3 Strain compatibility matrix among monokaryon strains by the hybridization experiment

Mycelium plugs (5 mm × 5 mm) of the tested strains were inoculated on petri dishes (60 mm diameter) containing 5 mL of PDA medium, with the plugs spaced 10– 15 mm apart. The cultures were incubated at 25°C for 10–12 days. After the mycelial plugs grew and contacted each other, the presence of clamp connections in the mycelia of the resulting colonies was observed using an upright microscope. If clamp connections were identified, the paired monokaryon strains were considered mating-type compatible; conversely, if any clamp connections were not observed, the two monokaryon strains were considered mating-type incompatible(Krings et al., 2011). The strain compatibility matrix among monokaryon strains was constructed from the clamp connections observation.

#### 2.2.4 A/B Loci consistency table for incompatible monokaryon pairs using the OWE-SOJ experiment

Monokaryon strains exhibiting incompatibility in single-cross experiments were inoculated onto PDA medium and incubated at 25°C for 6 days. Subsequently, a 5 mm diameter mycelium plug from the edge of a purified colony of two mutually incompatible strains was taken using a puncher and inoculated onto an OWE medium, with the plugs spaced 5 mm apart. After the mycelia of the paired strains began to contact, an agar strip approximately 2 mm wide and 35 mm long containing the two inoculation blocks was cut perpendicular to the interface line and transferred to a SOJ medium. The cultures were incubated at 25°C for 4-6 days, and the colonies phenotype was recorded. Referencing the results of mating-type analysis using the OWE-SOJ experiment of *Lentinula edodes*, the A/B loci consistency table was determined based on colony characteristics(Au et al., 2014; Fangcan et al., 2000; Yu et al., 2022). Strains forming a velvety colony along the agar strip without clamp connections exhibited an A=B≠ mating reaction. Conversely, strains forming a sparse hyphal contact zone between the two inoculation blocks’ colonies without clamp connections exhibited an A≠B= mating reaction.

## 3. Results

### 3.1 Implementation of MTI for mating-type inference

The fundamental principle of mating-type inference in MTI is based on the exhaustive enumeration method. This method involves directly traversing all possible mating-types among strains, generating the whole space of mating-type combinations. By calculating the score for all combinations to represent their coherence with the strain compatibility matrix and A/B loci consistency table, the combination with the highest score is recognized as the most likely monokaryon mating-types. However, the number of combinations exponentially increases with the population size of monokaryons. For example, 10 monokaryons could lead to 10^20^ possible mating-type combinations, making the score computation for those combinations is hardly finished in a reasonable time. To overcome the problem, MTI employs the combinatorial pruning traversal algorithm by taking advantage of the high mating compatibility rate among monokaryons. The basic logic of the algorithm is that fixing monokaryons with clearly defined mating-types to reduce the combination space among strains. To this end, the specific operation steps are as follows. Firstly, sub-population with strains that are pairwise compatible in the strain compatibility table are selected, since the mating-types of these strains could be easily determined. Secondly, the exhaustive enumeration method is used to generate all possible mating-type combinations of remaining monokaryons, thereby significantly reducing the complexity of score calculations. Taking a population of 30 monokaryons with 90% mating compatibility rate as an example, the combinatorial pruning traversal algorithm can reduce the 27,000 possible mating-type combinations to 6 ones (**Figure 3**), significantly improving the efficiency of mating-type inference.

**Figure 3:**
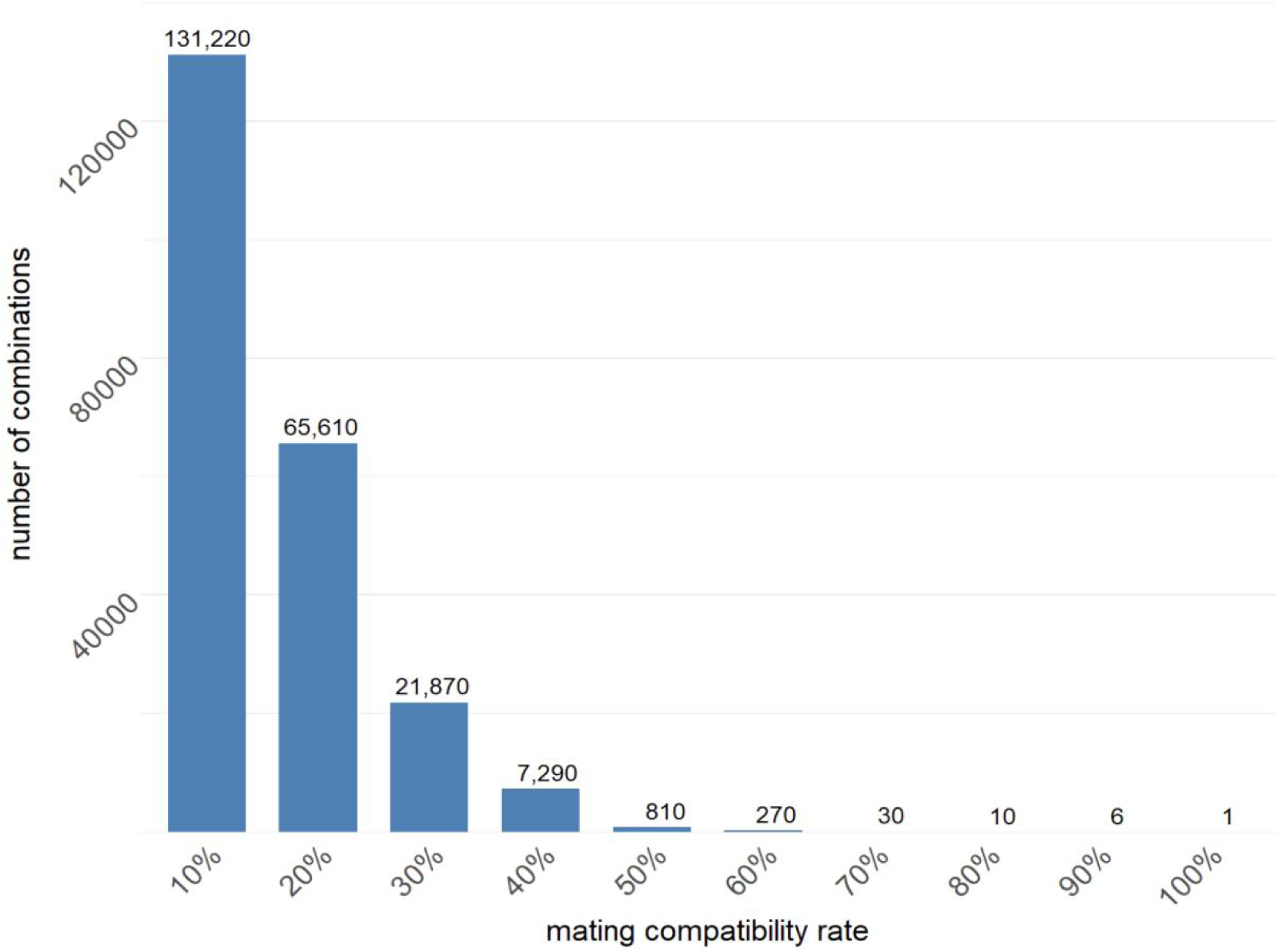
Number of traversing combinations with various mating compatibility rate. The number of mating-type combinations were listed on the bar.

### 3.2 Application platform developed with MTI

The source code for MTI software has now been uploaded to the GitHub platform (https://github.com/bxx2004/MTI-web). The example data and usage documents of the software are included in the GitHub repository. To facilitate the application of this software, we have developed an online web platform for tetrapolar basidiomycete mating-type inference based on MTI. This platform consists of three modules: data input, statistics and analysis, and result display (**Figure 4**). The data input module is used for user data upload, the statistics and analysis module is used for analysis and inference based on input data, and the result display module is used for displaying and printing mating-type results.

**Figure 4:**
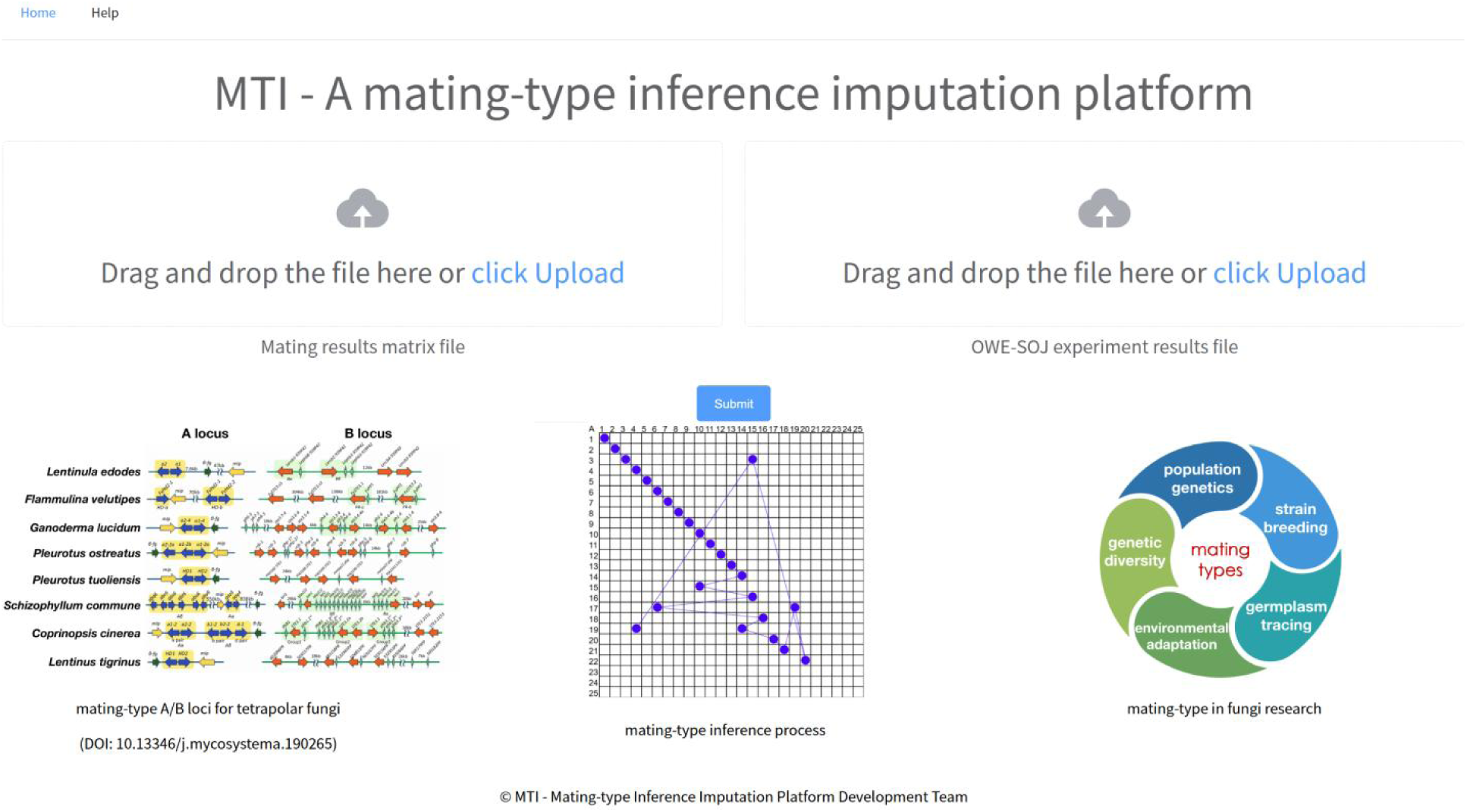
The snapshot of online web platform with MTI. The online application needs the input of monokaryon parent table, strain compatibility matrix and A/B loci consistency table.

In the data input module, users can upload data tables by clicking. To ensure the accuracy and reproducibility of mating-type inference, MTI requires users to provide the following standardized input files (**Table 2**): 1) monokaryon parent table, including the diploid parentage information of monokaryon strains; 2) strain compatibility matrix, including results of clamp connection observation in monokaryon hybridization experiments; 3) A/B loci consistency table, including results of colony trait observations from OWE-SOJ experiment for all incompatible hybridizations. The MTI platform provides example data and detailed file format descriptions. After submitting the data, users can click the “Run” button to perform mating-type inference.

**Table 2:**
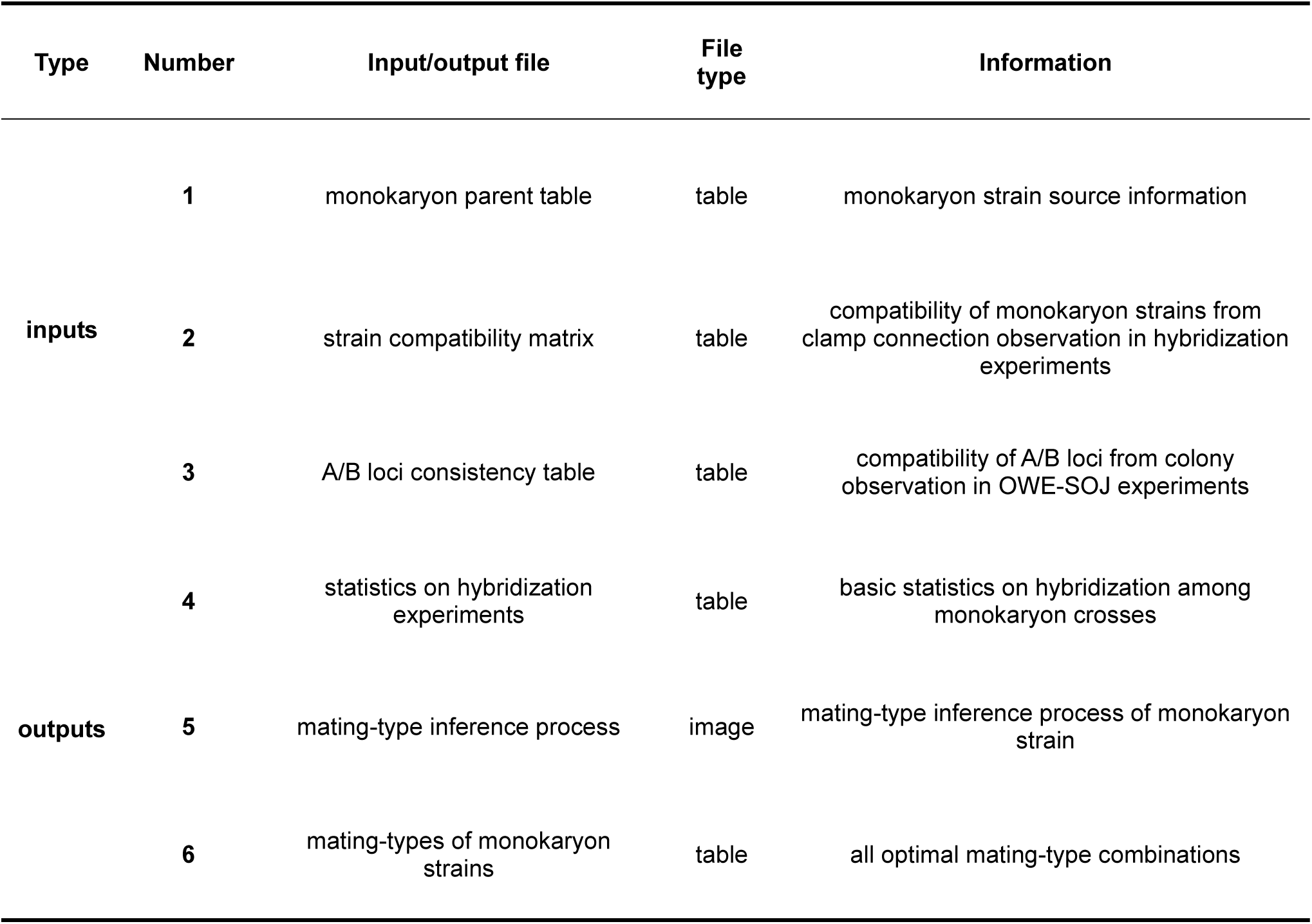
Summary of Input and Output Results for MTI.

| Type | Number | Input/output file | File type | Information |
| --- | --- | --- | --- | --- |
| inputs | 1 | monokaryon parent table | table | monokaryon strain source information |
|  | 2 | strain compatibility matrix | table | compatibility of monokaryon strains from clamp connection observation in hybridization experiments |
|  | 3 | A/B loci consistency table | table | compatibility of A/B loci from colony observation in OWE-SOJ experiments |
|  | 4 | statistics on hybridization experiments | table | basic statistics on hybridization among monokaryon crosses |
| outputs | 5 | mating-type inference process | image | mating-type inference process of monokaryon strain |
|  | 6 | mating-types of monokaryon strains | table | all optimal mating-type combinations |

The statistics and analysis module is the core processing module of the program, carrying out statistical analysis of the input data, performing the combinatorial pruning traversal algorithm for mating-type inference and output result for the next results display module (**Table 2**).

The results display module concentrates on visualization of the mating-type inference process, and mating-type result statistics (**Table 2**). The hybridization results statistics section lists the basic information such as the number of monokaryon strains, the number of hybridization experiments and missing hybridization combinations from the input data. The mating-type matrix is used to visualize the mating-type inference process for monokaryon strains. The mating-type result statistics employs charts to illustrate the distribution of mating-types across all monokaryon strains, along with the specific strain information associated with each mating-type.

### 3.3 Mating-type inference for *Flammulina velutipes* using MTI

To illuminate the application of the software developed in this work, mating-types of *Flammulina velutipes* were inferred with MTI using results of hybridization experiments. To this end, 30 monokaryon strains were subjected to pairwise hybridization experiments, yielding a total of 420 hybridization results. Based on the microscopic examination of clamp connection, 410 and 10 pairs showed compatibility and incompatible, respectively (**Supplementary Table 1**). 10 incompatible hybridization pairs were subjected to OWE-SOJ experimental results to probe the consistency of A/B loci (**Supplementary Table 2**).

Through the aforementioned experiments, the input file obtained from 30 *Flammulina velutipes* monokaryon strains hybridization experiments was uploaded to the online platform of MTI. Upon submitting the analysis, the system returned the statistics of input data and performed mating-type inference of 30 monokaryon strains (**Figure 5**). As a result, MTI inferred the best mating-type combination for the 30 monokaryon strains with the highest score of 870 to satisfy all hybridization experiment observations, with 24 alleles at the A locus and 27 alleles at the B locus (**Figure 6a**).

**Figure 5:**
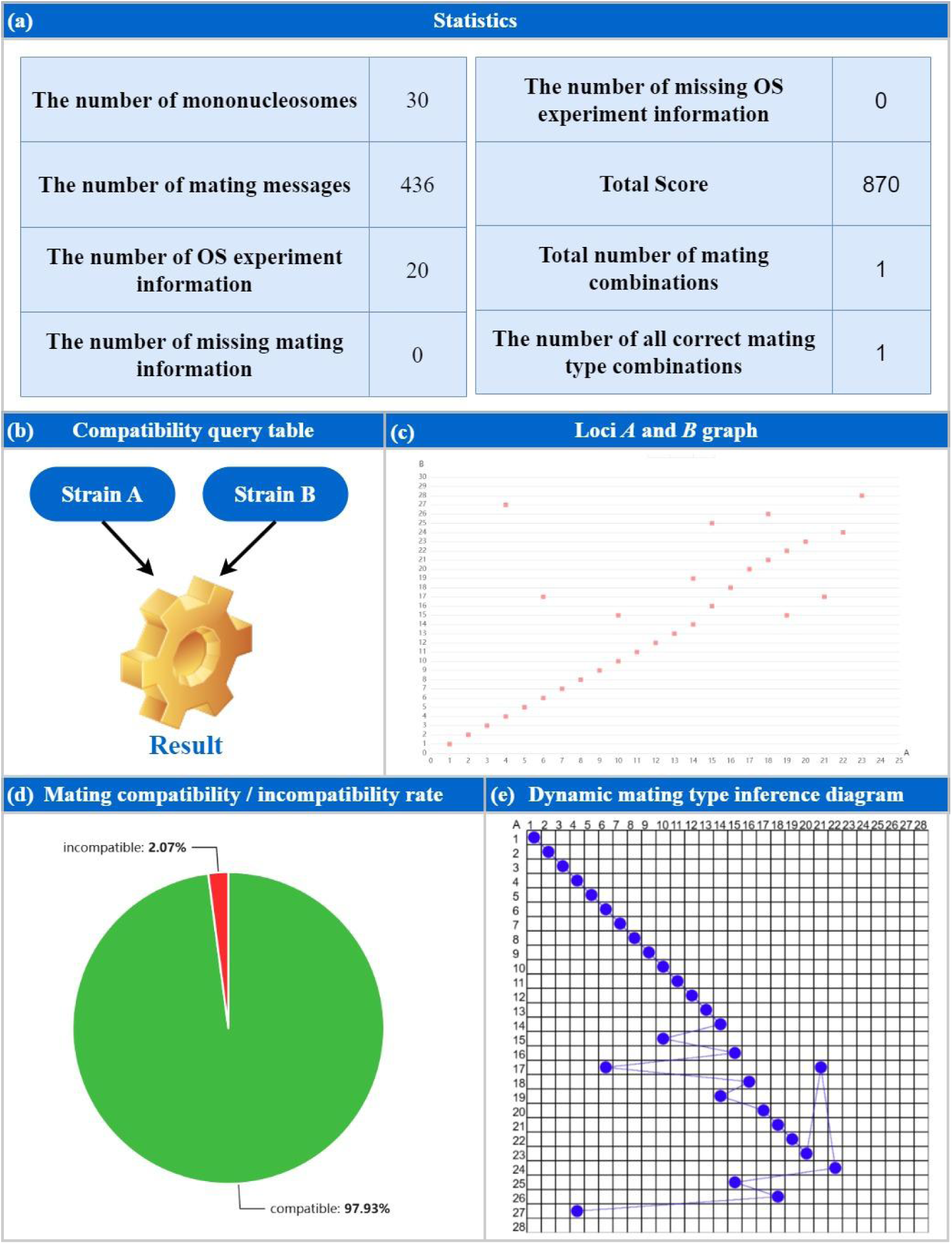
The results of mating-type inference using MTI platform for *Flammulina velutipes*. (a) The statistics for input data and mating-type inference; (b) compatibility query function for strains; (c) The distribution of mating-types for all strains; (d) compatibility and incompatibility rate among strains; (e) Dynamic diagram for mating-type inference.

**Figure 6:**
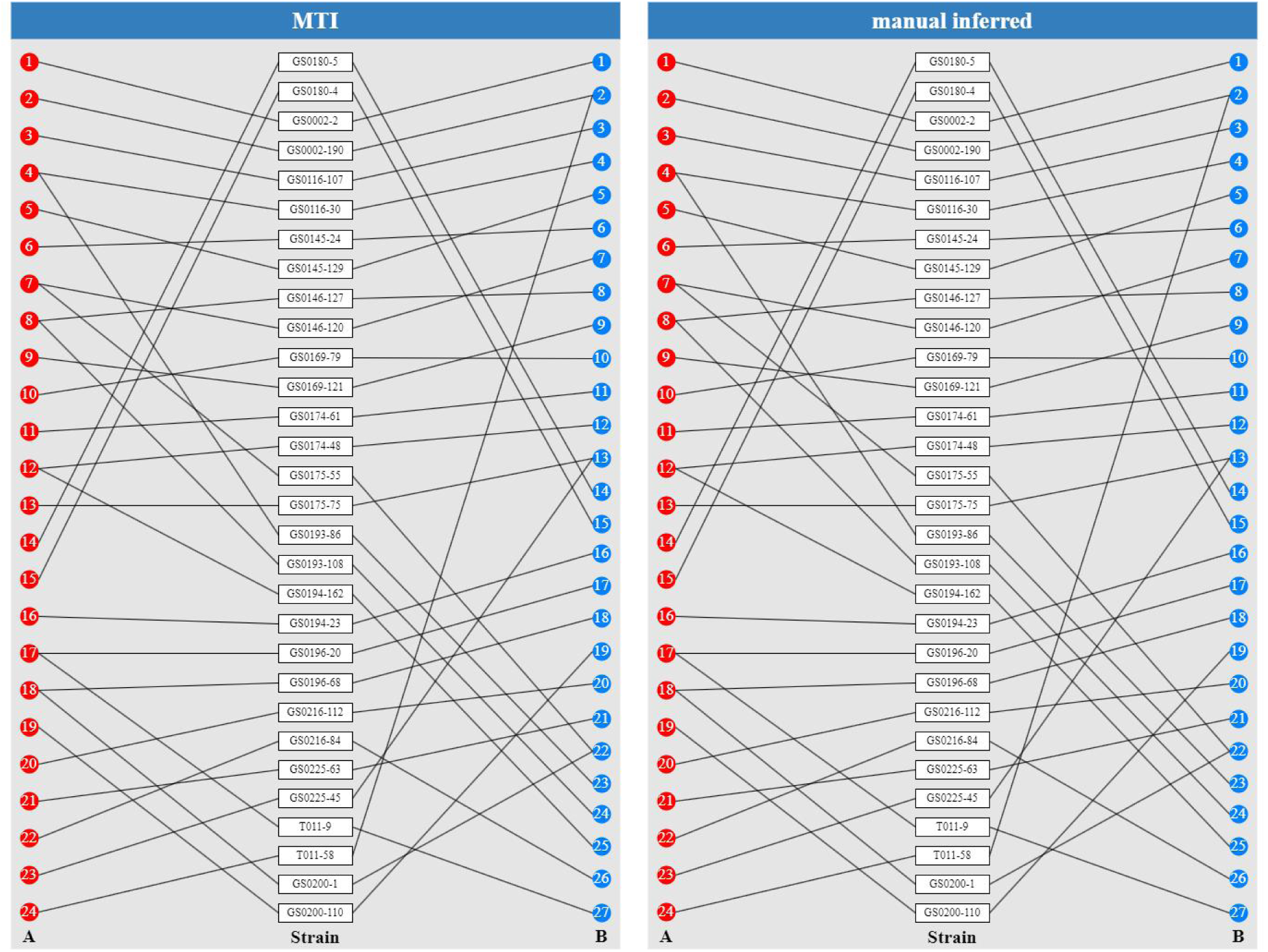
The validation of MTI by comparing with the manual inferred result. The mating-type combination from MTI (left) and manual inferred (right) result are identical.

### 3.4 Manual mating-type inference to validate MTI result

To validate the result of mating-type inference by MTI, we also performed manual investigation to determine the mating-types of 30 monokaryon strains (**Supplementary Note 1**). According to the strain compatibility matrix and A/B loci consistency table from hybridization and OWE-SOJ experiments (**Supplementary** Figure 1), the mating-types of those 30 monokaryons were inferred manually, and the detailed processes were recorded in the **Supplementary Note 1**. The whole manual inference takes about 3 days to draw the conclusion. As a result, the manually induced mating-types of the 30 monokaryons were identical to MTI results (**Figure 6**), validating mating-type inference of the software.

### 3.5 How errors in hybridization experiments influence mating-type inference results

Hybridization experiments provide the basic information for mating-type inference(Wu et al., 2013); therefore, errors in the strain compatibility matrix pose potential challenges for accurate mating-type determination. However, how errors in hybridization experiments influence mating-type inference is still obscure. MTI enables us to investigate how input influences the result of mating-type inference using simulated data. False-positive and false-negative are two common errors in hybridization experiments. To simulate hybridization experiments with errors, we randomly introduced 1 to 5 spurious compatible (false-positive) or incompatible (false-negative) records in the strain compatibility matrix among 30 monokaryons. The mating-type of 30 monokaryons were inferred using simulated datasets with MTI and compared with the true counterpart.

Our results revealed the distinct effects of false-positives and false-negatives on mating-type inferences. Any false-positive in the strain compatibility matrix could mislead the MTI to draw a wrong best mating-type combination, although the combination could achieve the full score using the wrong simulated dataset (**Figure 7a and 7b**). In contrast, although false-negatives in the simulated strain compatibility matrix could lower the mating-type combination score; however, if there is only one false-negative, the resulting best mating-type combination is identical to the true combination, but when the false-negative number is larger than 1, many alternative best mating-type combinations could also be inferred, including the true one (**Figure 7c**).

**Figure 7:**
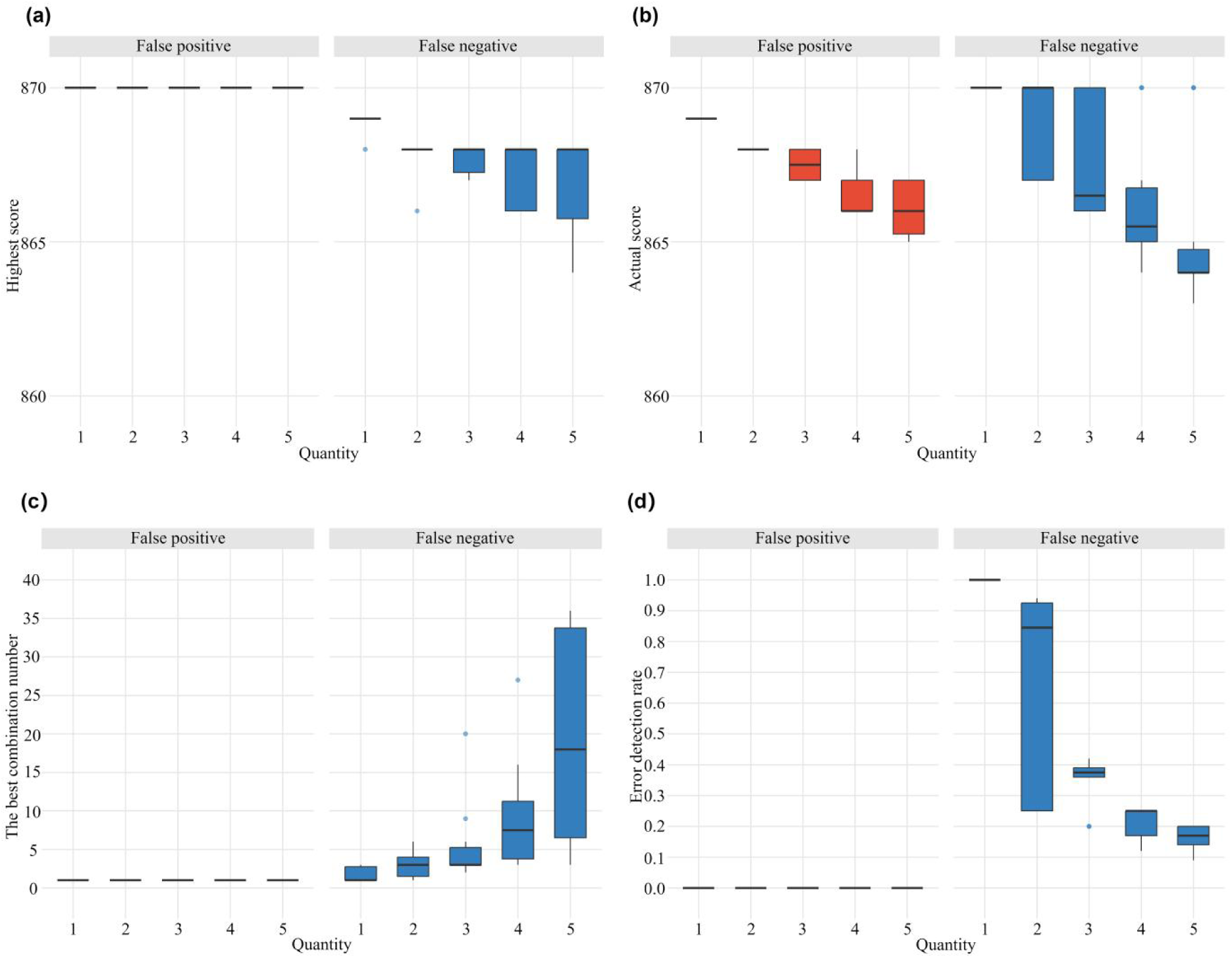
The impact of false-positives and false-negatives on mating-type inference. (a) The effect of random false-positives and false-negatives on the highest score. (b) The effect of random false-positives and false-negatives on the actual score. (c) The effect of random false-positives and false-negatives on the optimal combination number. (d) The effect of random false-positives and false-negatives on the accuracy of false-positive detection.

Since errors in hybridization experiments could mislead the mating-type inference, we then investigated if MTI could be used to detect false-positives or false-negatives records in the strain compatibility matrix. From our analysis, if the number of false-negatives was small, especially in the case of 1 false-negative, MTI could accurately detect the error in the compatibility matrix, providing valuable information for experimental verification and tracing (**Figure 7d**). However, MTI failed to detect any false-positives in our simulation analysis, since false-positives could always lead to the full-scored mating-type combination and satisfy the simulated strain compatibility matrix(**Figure 7d**). Our simulation analysis indicated that false-positives exert more substantial influence on mating-type inference and were difficult to be traced. Therefore, greater emphasis should be placed to avoid false-positive records during hybridization experiments, and the criteria for hybridization compatibility recognition should be more stringent in practice.

## 4. Discussions

### 4.1 MTI as the first mating-type inference tool for tetrapolar fungi

The vast majority of medicinal and edible fungi are basidiomycetes and generally exhibit high mating-type diversity(Coelho et al., 2017). Mating-type identification serves as the foundation for studying the genetic diversity and breeding of those species(Ha et al., 2018). Currently, the basic method for mating-types identification relies on monokaryon hybridization experiments(A. Li et al., 2015). The manual mating-types inference from the strain compatibility matrix and A/B loci consistency is exceptionally labor-intensive and error-prone for large scale monokaryon populations. This study established an automatic, rapid and accurate method for inferring mating-types in A/B loci of tetrapolar fungi monokaryon strains using the strain compatibility matrix from clamp connection observations in hybridization experiments(Wu et al., 2013) and A/B loci consistency table from colony phenotype records in OWE-SOJ experiments(Ryan & Smith, 2004). MTI were then further applied and validated by the *F. velutipes* monokaryon population. Specifically, the mating-types of 30 monokaryon strains were rapidly and accurately inferred by MTI and were identical with manual inferred results. In summary, MTI provides the first mating-type inference tool for the majority of edible and medicinal fungi, as well as other tetrapolar fungi.

### 4.2 Main features of MTI for mating-type inference

Addressing the task of large-scale mating-type identification, the mating-type inference tool developed in this study for tetrapolar basidiomycetes possesses the following features in automation, speed, accuracy and intelligence. Firstly, MTI is automatic for the mating-type inference, since MTI could infer mating-type without requiring any manual logical reasoning, significantly lowering the threshold and difficulty associated with large-scale mating-type inference. Secondly, MTI is fast for the mating-type inference, especially in large populations. In the case of 30 *F. velutipes* monokaryons, MTI required only two minutes to generate the result, but the manual inference with the identical input would take several days for experienced researchers. Thirdly, MTI is accurate, which is validated by the *F. velutipes* population. MTI relies on programmatic logic for mating-type inference, avoiding subjective errors in the manual manner. Fourthly, MTI is smart, since the software not only infers the best mating-type combination for monokaryon strains, but also could identify potential errors in the strain compatibility matrix, providing valuable information for experimental design and tracing.

### 4.3 Technical scheme for mating-type identification for tetrapolar fungi using MTI

Based on the features and functions of MTI, we propose a standard technical scheme of monokaryon mating-types inference and mating-type library construction for tetrapolar fungi, including the majority of edible and medicinal fungi, which mainly includes the following steps:

1) Protoplast monokaryons are isolated from dikaryon parents, and the monokaryon parent table is obtained;
2) Pair-wise hybridization experiments of monokaryons are performed and the strain compatibility matrix is obtained from the clamp connection observation;
3) OWE-SOJ experiments are carried out for incompatible monokaryon pairs and the A/B loci consistency table is obtained from colony phenotype observation;
4) Using the monokaryon parent table, the strain compatibility matrix and the A/B loci consistency table, the mating-types of all monokaryons are inferred using MTI software, which is used to establish the mating-type library of the species with the representative strains;
5) For a new monokaryon with unknown mating-type, hybridization experiments are performed for the new monokaryon and representative strains in the mating-type library;
6) The mating-type of the new monokaryon is determined with MTI software using the hybridization experiment results. If the new monokaryon possesses an identical mating-type in the library, add the monokaryon to the strain list of this mating-type. Otherwise, if the new monokaryon provides a new mating-type, then the mating-type library is updated and the new monokaryon is listed as the representative strain for the new mating-type.

## 5. Conclusion

The MTI software provides an automatic, rapid, accurate and smart solution for mating-types inference of tetrapolar fungi, especially for large-scale populations. As the first tools specific for mating-types inference, MTI not only simplifies the process of mating-type identification of tetrapolar fungi, but also lays a solid foundation for a standardized research paradigm for the mating-type library construction for those species, which plays an important role in fungal genetic diversity, germplasm tracing and strain breeding(Breene, 1990; Lee et al., 2010). Therefore, MTI offers an efficient analytical tool for the field of fungal genetics and diversity, holding significant theoretical value and promising applications(Tang et al., 2016).

## Acknowledgement

This work was financed by the National Natural Science Foundation of China (No. 32270028) and the National key R&D Program of China (2023YFF1000800).

## Data Availability Statement

The software and data have been uploaded to the GitHub (https://github.com/bxx2004/MTI-web) and are accessible via the MTI online platform (http://124.222.225.24:82/).

## Author Contributions

Dapeng Bao and Shijun Xiao conceptualized the study and designed the methodology. Haixu Liu developed the program and online platform. Yu Wang conducted the laboratory work. Weijun Li, Huiyang Xiong, and Ruiheng Yang performed feasibility validation of the program. All authors read then approved the manuscript for publication.

