## Supplementary Note for "Mating-Type Imputation (MTI) provides an efficient tool for the mating-type inference of tetrapolar fungi"

**The detailed mating-type manual inference for 30 *Flammulina velutipes* monokaryons**

Based on the hybridization results, the 30 tested strains could be divided into two groups. The first group consisted of 21 strains that exhibited pairwise compatibility within the group, resulting in 410 compatible hybridization outcomes. These 21 tested strains were specifically GS0002-2, GS0002-190, GS0116-107, GS0116-30, GS0145-129, GS0145-24, GS0146-120, GS0146-127, GS0169-121, GS0169-79, GS0174-61, GS0174-48, GS0175-75, GS0180-4, GS0180-5, GS0194-23, GS0196-20, GS0196-68, GS0200-110, GS0216-112, and GS0225-63. The results of these compatible hybridizations indicated that the mating types of these strains were all different A≠B≠ combinations, each unique. Following this principle, the mating types of the strains in this group could be defined as $A_1B_1$, $A_2B_2$, $A_3B_3$, ......, $A_{21}B_{21}$, i.e., GS0002-2 ($A_1B_1$), GS0002-190 ($A_2B_2$), GS0116-107 ($A_3B_3$), GS0116-30 ($A_4B_4$), GS0145-129 ($A_5B_5$), GS0145-24 ($A_6B_6$), GS0146-120 ($A_7B_7$), GS0146-127 ($A_8B_8$), GS0169-121 ($A_9B_9$), GS0169-79 ($A_{10}B_{10}$), GS0174-61 ($A_{11}B_{11}$), GS0174-48 ($A_{12}B_{12}$), GS0175-75 ($A_{13}B_{13}$), GS0180-4 ($A_{14}B_{14}$), GS0180-5 ($A_{15}B_{15}$), GS0194-23 ($A_{16}B_{16}$), GS0196-20 ($A_{17}B_{17}$), GS0196-68 ($A_{18}B_{18}$), GS0200-110 ($A_{19}B_{19}$), GS0216-112 ($A_{20}B_{20}$), and GS0225-63 ($A_{21}B_{21}$).

For further analysis, based on the hybridization results, the 30 tested strains could be divided into two groups. The first group consisted of 21 strains that exhibited pairwise compatibility within the group, resulting in 210 compatible hybridization outcomes. These 21 tested strains were specifically GS0002-2, GS0002-190, GS0116-107, GS0116-30, GS0145-129, GS0145-24, GS0146-120, GS0146-127, GS0169-121, GS0169-79, GS0174-61, GS0174-48, GS0175-75, GS0180-4, GS0180-5, GS0194-23, GS0196-20, GS0196-68, GS0200-110, GS0216-112, and GS0225-63. The results of these compatible hybridizations indicated that the mating-types of these strains were all different A≠B≠ combinations, each unique. Following this principle, the mating-types of the strains in this group could be defined as $A_1B_1$, $A_2B_2$, $A_3B_3$, ......, $A_{21}B_{21}$, i.e., GS0002-2 ($A_1B_1$), GS0002-190 ($A_2B_2$), GS0116-107 ($A_3B_3$), GS0116-30 ($A_4B_4$), GS0145-129 ($A_5B_5$), GS0145-24 ($A_6B_6$), GS0146-120 ($A_7B_7$), GS0146-127 ($A_8B_8$), GS0169-121 ($A_9B_9$), GS0169-79 ($A_{10}B_{10}$), GS0174-61 ($A_{11}B_{11}$), GS0174-48 ($A_{12}B_{12}$), GS0175-75 ($A_{13}B_{13}$), GS0180-4

($A_{14}B_{14}$), GS0180-5 ($A_{15}B_{15}$), GS0194-23 ($A_{16}B_{16}$), GS0196-20 ($A_{17}B_{17}$), GS0196-68 ($A_{18}B_{18}$), GS0200-110 ($A_{19}B_{19}$), GS0216-112 ($A_{20}B_{20}$), and GS0225-63 ($A_{21}B_{21}$).

The second group included the remaining 9 strains. Among these 9 strains, 36 hybridization pairs were produced, of which 34 were compatible and 2 were incompatible, specifically between GS0175-55 and GS0200-1, and between GS0193-86 and GS0216-84. Additionally, these 9 strains could form 189 hybridization combinations with the aforementioned 21 strains, resulting in 181 compatible pairs and 8 incompatible pairs. The incompatible pairs were specifically between GS0175-55 and GS0146-120, GS0193-86 and GS0116-30, GS0193-108 and GS0146-127, GS0194-162 and GS0174-48, GS0200-1 and GS0196-68, GS0225-45 and GS0175-75, T011-9 and GS0196-20, and T011-58 and GS0002-190 (**Supplementary Table 1**).

**Supplementary Table 1: hybridization experiment results among 9 strains with undetermined mating-types.**

| Strain number | GS0175-55 | GS0193-86 | GS0193-108 | GS0194-162 | GS0216-84 | GS0225-45 | T011-9 | T011-58 | GS0200-1 |
|---|---|---|---|---|---|---|---|---|---|
| GS0002-2 | ＋ | ＋ | ＋ | ＋ | ＋ | ＋ | ＋ | ＋ | ＋ |
| GS0002-190 | ＋ | ＋ | ＋ | ＋ | ＋ | ＋ | ＋ | － | ＋ |
| GS0116-107 | ＋ | ＋ | ＋ | ＋ | ＋ | ＋ | ＋ | ＋ | ＋ |
| GS0116-30 | ＋ | － | ＋ | ＋ | ＋ | ＋ | ＋ | ＋ | ＋ |
| GS0145-24 | ＋ | ＋ | ＋ | ＋ | ＋ | ＋ | ＋ | ＋ | ＋ |
| GS0145-129 | ＋ | ＋ | ＋ | ＋ | ＋ | ＋ | ＋ | ＋ | ＋ |
| GS0146-127 | ＋ | ＋ | － | ＋ | ＋ | ＋ | ＋ | ＋ | ＋ |
| GS0146-120 | － | ＋ | ＋ | ＋ | ＋ | ＋ | ＋ | ＋ | ＋ |
| GS0169-79 | ＋ | ＋ | ＋ | ＋ | ＋ | ＋ | ＋ | ＋ | ＋ |
| GS0169-121 | ＋ | ＋ | ＋ | ＋ | ＋ | ＋ | ＋ | ＋ | ＋ |
| GS0174-61 | ＋ | ＋ | ＋ | ＋ | ＋ | ＋ | ＋ | ＋ | ＋ |
| GS0174-48 | ＋ | ＋ | ＋ | － | ＋ | ＋ | ＋ | ＋ | ＋ |
| GS0175-55 | \ | ＋ | ＋ | ＋ | ＋ | ＋ | ＋ | ＋ | － |
| GS0175-75 | ＋ | ＋ | ＋ | ＋ | ＋ | － | ＋ | ＋ | ＋ |
| GS0193-86 | ＋ | \ | ＋ | ＋ | － | ＋ | ＋ | ＋ | ＋ |
| GS0193-108 | ＋ | ＋ | \ | ＋ | ＋ | ＋ | ＋ | ＋ | ＋ |
| GS0194-162 | ＋ | ＋ | ＋ | \ | ＋ | ＋ | ＋ | ＋ | ＋ |
| GS0194-23 | ＋ | ＋ | ＋ | ＋ | ＋ | ＋ | ＋ | ＋ | ＋ |
| GS0196-20 | ＋ | ＋ | ＋ | ＋ | ＋ | ＋ | － | ＋ | ＋ |
| GS0196-68 | ＋ | ＋ | ＋ | ＋ | ＋ | ＋ | ＋ | ＋ | － |
| GS0216-112 | ＋ | ＋ | ＋ | ＋ | ＋ | ＋ | ＋ | ＋ | ＋ |
| GS0216-84 | ＋ | － | ＋ | ＋ | \ | ＋ | ＋ | ＋ | ＋ |
| GS0225-63 | ＋ | ＋ | ＋ | ＋ | ＋ | ＋ | ＋ | ＋ | ＋ |
| GS0225- | ＋ | ＋ | ＋ | ＋ | ＋ | \ | ＋ | ＋ | ＋ |

| | | | | | | | | | |
|---|---|---|---|---|---|---|---|---|---|
| 45 | | | | | | | | | |
| T011-9 | + | + | + | + | + | + | \ | + | + |
| T011-58 | + | + | + | + | + | + | + | \ | + |
| GS0200-1 | − | + | + | + | + | + | + | + | \ |
| GS0200-110 | + | + | + | + | + | + | + | + | + |
| GS0180-4 | + | + | + | + | + | + | + | + | + |
| GS0180-5 | + | + | + | + | + | + | + | + | + |

These 9 strains exhibited incompatible hybridization results, necessitating further inference of their mating-types. Until their mating-types are clearly determined, they are all denoted as $A_xB_x$, where x represents any possible index number. The incompatible hybridization between same mating-type, these two monokaryon strains would exhibit identical hybridization results when crossed with other monokaryon strains.

Among the 9 strains in the second group, 3 monokaryon strains (GS0175-55, GS0193-86, and GS0200-1) exhibited two incompatible hybridization results. GS0175-55 showed incompatibility when hybridized with GS0146-120 ($A_7B_7$) and GS0200-1 ($A_xB_x$). Since GS0146-120 ($A_7B_7$) and GS0200-1 ($A_xB_x$) were compatible, their mating-types were completely different, suggesting that the mating-type of GS0175-55 could not be of the A=B= type. Its possible mating-types were ($A_7B_x$) or ($A_xB_7$). Given that GS0175-55 was compatible with all other tested Monokaryon strains, the x in its mating-type ($A_7B_x$) or ($A_xB_7$) represented a new mating-type, which could be named 22. Thus, the mating-type of GS0175-55 was either ($A_7B_{22}$) or ($A_{22}B_7$).

GS0193-86 showed incompatibility when hybridized with GS0116-30 ($A_4B_4$) and GS0216-84 ($A_xB_x$). Since GS0116-30 ($A_4B_4$) and GS0216-84 ($A_xB_x$) were compatible, their mating-types were completely different, suggesting that the mating-type of GS0193-86 could not be of the A=B= type. Its possible mating-types were ($A_4B_x$) or ($A_xB_4$). Given that GS0193-86 was compatible with all other tested Monokaryon strains, the x in its mating-type ($A_4B_x$) or ($A_xB_4$) represented a new mating-type, which could be named 23. Thus, the mating-type of GS0193-86 was either ($A_4B_{23}$) or ($A_{23}B_4$).

If the mating-type of GS0175-55 was ($A_7B_{22}$), then the mating-type of GS0200-1 would be ($A_{18}B_{22}$), or if the mating-type of GS0175-55 was ($A_{22}B_7$), then the mating-type of GS0200-1 would be ($A_{22}B_{18}$).

Among the 9 strains in the second group, 6 Monokaryon strains (GS0193-108, GS0194-162, GS0216-84, GS0225-45, T011-9, and T011-58) exhibited one incompatible hybridization result.

GS0193-108 showed incompatibility when hybridized with GS0146-127 (A8B8), suggesting that its mating-type could be one of ($A_8B_8$), ($A_8B_x$), or ($A_xB_8$). Since GS0193-108 was compatible with all other Monokaryon strains, the x in its mating-type ($A_8B_x$) or ($A_xB_8$) represented a new type, which in this study was assigned the serial number 24. Thus, the mating-type of GS0193-108 was either ($A_8B_8$), ($A_8B_{24}$), or ($A_{24}B_8$).

By analogy, the mating-type of GS0194-162 could be ($A_{12}B_{12}$), ($A_{12}B_{25}$), or ($A_{25}B_{12}$); the mating-type of GS0216-84 could be ($A_{26}B_{23}$) or ($A_{23}B_{26}$) (since the compatibility performance of GS0216-84 differed from that of GS0193-86, the A=B= possibility was excluded); the mating-type of GS0225-45 could be ($A_{13}B_{13}$), ($A_{13}B_{27}$), or ($A_{27}B_{13}$); the mating-type of T011-9 could be ($A_{17}B_{17}$), ($A_{17}B_{28}$), or ($A_{28}B_{17}$). The mating-type of T011-58 could be ($A_{29}B_2$), ($A_2B_{29}$), or ($A_{29}B_2$).

Further inference of the A and B mating-type information for these 9 strains requires clarification based on the OWE-SOJ experimental results.

### 3.4 Results of the OWE-SOJ Experiment

In the OWE-SOJ experiment, the mating-type relationship between the strains in different hybridization combinations is determined based on the colony characteristics, specifically whether it is A=B≠ or A≠B=, thereby further clarifying the mating-type of the strains under investigation (**Supplementary Table 2**). The phenotypic characteristics of the OWE-SOJ experimental results are illustrated in **Supplementary Figure 1**.

**Supplementary Table 2: OWE-SOJ experiment results and mating-type analysis of 9 strains with undetermined mating-types.**

| Strains with undetermined mating-types. | Hybrid pairing strains and mating-types. | Phenotype of the OWE-SOJ experiment. | Types of results for the OWE-SOJ experiment. | Putative mating-type. |
|---|---|---|---|---|

| | | | | |
|---|---|---|---|---|
| GS0175-55（*A7B22* or *A22B7*） | GS0146-120（*A7B7*） | Cottony colonies without clamp connections. | *A*＝*B*≠ | *A7B22* |
| | GS0200-1（*A18B22* or *A22B18*） | A contact zone composed of sparse hyphae without clamp connections. | *A*≠*B*＝ | *A7B22* |
| GS0193-86（*A4B23* or *A23B4*） | GS0116-30（*A4B4*） | Cottony colonies without clamp connections. | *A*＝*B*≠ | *A4B23* |
| | GS0216-84（*A26B23* or *A23B26*） | A contact zone composed of sparse hyphae without clamp connections. | *A*≠*B*＝ | *A4B23* |
| GS0200-1（*A18B22* or *A22B18*） | GS0175-55（*A7B22 or A22B7*） | A contact zone composed of sparse hyphae without clamp connections. | *A*≠*B*＝ | *A18B22* |
| | GS0196-68（*A18B18*） | Cottony colonies without clamp connections. | *A*＝*B*≠ | *A18B22* |
| GS0193-108（*A8B8* or *A8B24* or *A24B8*） | GS0146-127（*A8B8*） | Cottony colonies without clamp connections. | *A*＝*B*≠ | *A8B24* |
| GS0194-162（*A12B12* or *A12B25* or *A25B12*） | GS0174-48（*A12B12*） | Cottony colonies without clamp connections. | *A*＝*B*≠ | *A12B25* |
| GS0216-84（*A26B23* or *A23B26*） | GS0193-86（*A4B23 or A23B4*） | Cottony colonies without clamp connections. | *A*＝*B*≠ | *A23B26* |
| GS0225-45（*A13B13* or *A13B27* or *A27B13*） | GS0175-75（*A13B13*） | A contact zone composed of sparse hyphae without clamp connections. | *A*≠*B*＝ | *A27B13* |
| T011-9（*A17B17* or *A17B28* or *A28B17*） | GS0196-20（*A17B17*） | Cottony colonies without clamp connections. | *A*＝*B*≠ | *A17B28* |
| T011-58（*A29B2* or *A2B29* or *A29B2*） | GS0002-190（*A2B2*） | Cottony colonies without clamp connections. | *A*＝*B*≠ | *A29B2* |

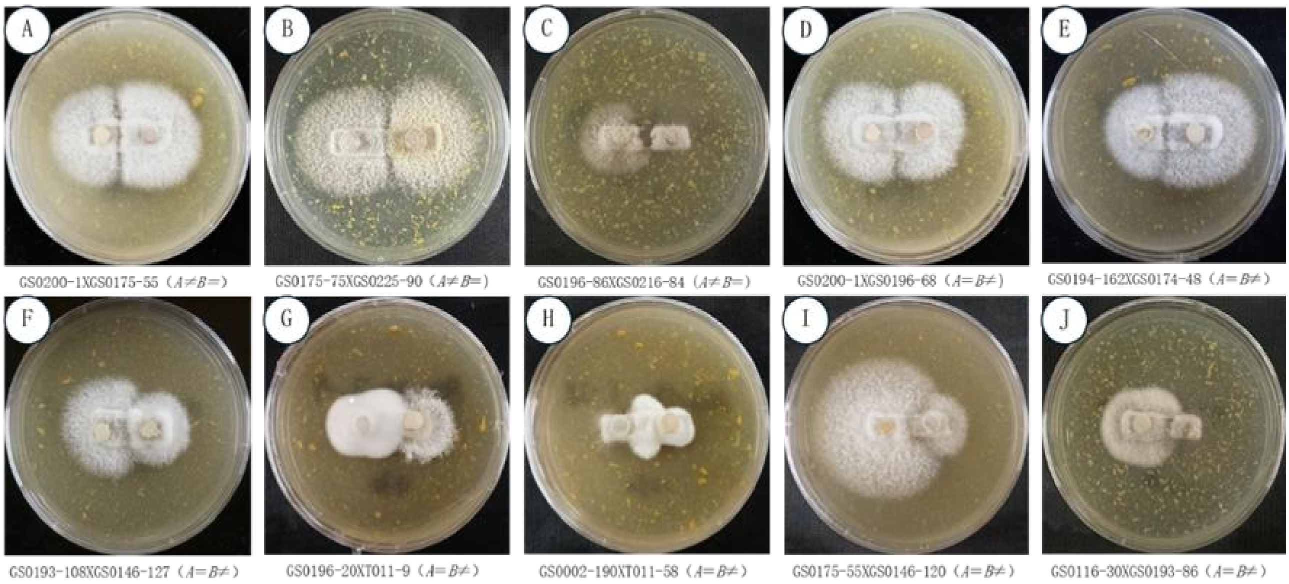


**Supplementary Figure 1: OWE-SOJ experiment for 9 strains with undetermined mating-types**
